# 2D-QSAR and Molecular Docking based virtual screening of the herbal molecules against Alzheimer’s Disorder: An approach to predict CNS activity

**DOI:** 10.1101/2022.10.08.511422

**Authors:** Aman Thakur, Arun Parashar, Vivek Sharma, Ajay Kumar, Vineet Mehta

## Abstract

Acetylcholinesterase (AChE) is one of the key enzyme targets that have been used clinically for the management of Alzheimer’s Disorder (AD). Numerous reports in the literature predict and demonstrate *in-vitro*, and *in-silico* anticholinergic activity of synthetic and herbal molecules, however, the majority of them failed to reproduce the results in preclinical or clinical settings. To address these issues, we developed a 2D-QSAR model that could not only efficiently predict the AChE inhibitory activity of herbal molecules but also predicted their potential to cross BBB to exert their beneficial effects during AD. Applying this model, virtual screening of the herbal molecules was performed and amentoflavone, asiaticoside, astaxanthin, bahouside, biapigenin, glycyrrhizin, hyperforin, hypericin, and tocopherol were predicted as the most promising herbal molecules for inhibiting AChE. Results were validated through molecular docking studies against human AChE (PDB ID: 4EY7). To determine whether or not these molecules can cross BBB to inhibit AChE within the CNS for being beneficial for the management of AD, we determined a CNS PPO score, which was found in the range of 1 to 3.76. Overall, the best results were observed for amentoflavone and our results demonstrated a PIC_50_ value of 7.377 nM, molecular docking score of −11.5 kcal/mol, and CNS MPO score of 3.76. In conclusion, we successfully developed a reliable and efficient 2D-QSAR model and predicted amentoflavone to be the most promising molecule that could inhibit human AChE enzyme within the CNS and could prove beneficial for the management of AD.

## 1. Introduction

Alzheimer’s Disorder (AD) is one of the most pervasive age-related neurodegenerative disorders that is associated with progressive neurodegeneration, dementia, and cognitive decline. The available therapeutic strategies for the management of AD are focused on eliminating the symptoms of AD and none of them is capable of reversing or halting the progression of AD and neurodegeneration (Kumar et. al., 2021). Lack of complete understanding regarding the pathogenesis of Alzheimer’s disease is the major hindrance in the development of new therapeutic strategies for AD (Kumar et. al., 2021). In the past few decades, the research work in identifying novel therapeutic strategies for the management of AD is focused on some of the prominent pathways of AD pathogenesis that includes the upregulation of cholinesterase signaling in the brain (Ferreira-Vieira et. al., 2016). Data from the clinical settings suggest that the pathogenesis of AD is presented by the breakdown of both neurochemical and structural neural networks, which include the cholinergic system (Hampel et. al., 2018). Upregulating cholinergic signaling in the brain has produced some promising beneficial results during AD (Hampel et. al., 2018). This is supported by the findings that treating rodents with cholinergic agonists resulted in marked improvement in memory and impaired cognitive performance and treatment with cholinergic antagonists resulted in deterioration of cognitive and memory functions (Ferreira-Vieira et. al., 2016; Chen et. al., 2022). The available drug candidates for targeting cholinergic signaling in the brain are only able to provide relief from the AD symptoms, besides, the effect on AD progression and reversal of neurodegeneration remains a major issue (Kumar et. al., 2021).

The effect of potential drug candidates on AD is governed to a large extent by their potential to cross the blood-brain barrier (BBB) (Pardridge et. al., 2009). In the past decade, a few drug candidates have been identified for the management of AD through upregulating cholinergic signaling, however, the majority of the drugs either fail to cross BBB or are having limited access to the brain, which not just reduced their efficacy against AD, but also require the use of the drug at higher dose that leads to toxic effects (Banks et. al., 2012). It is also worth mentioning that no major new drug has been approved by the US Food and Drug Administration (FDA) since 2003. Focussing research on the development of promising anticholinesterase (AChE) inhibitors for the management of AD, with the potential to cross BBB, have acceptable side effects, and could halt or reverse the AD progression is the dire need of the hour.

Herbal molecules are better tolerated in the human body than synthetic molecules and therefore, the focus on developing new candidates for AD management has shifted to herbal molecules in recent times (Thomford et. al., 2018). It is a well-established fact that the traditional drug discovery process is a highly time-consuming and expensive affair. Screening herbal molecules against AD is another major concern due to the lack of appropriate tools or models. Computer Aided Drug Designing (CADD) is one of the promising solutions for reducing the preclinical trial time and cost by employing structure-based drug designing (SBDD) or Ligand-based drug designing (LBDD) approaches to predict potential molecules (Yu et. al., 2017; Surabhi and Singh, 2018). LBDD technique such as Quantitative Structural Activity Relationship (QSAR) calculates the structural properties of already established compounds against a particular disease and relates it to their biological activity (Yu et. al., 2017; Surabhi and Singh, 2018). The SBDD approach which employs the Molecular Docking technique predicts the active conformation of the ligand in the binding pocket of the receptor and calculates the energy of the ligand-receptor complex (Yu et. al., 2017; Surabhi and Singh, 2018). This not just reduces the investment in term of time and funds, but also provide us with promising lead molecules with a higher predicted success rate against disorders like AD.

In the present study, we are focused on developing a 2D-QSAR model from the reported 32 tacrine-cinnamic acid hybrid molecules and using this model for the screening of herbal molecules for AChEinhibitory activity. Moreover, the results of the developed model were validated through molecular docking studies and CNS MPO score studies to predict not just activity against AChE, but also to predict potential to cross BBB and act on the CNS.

## 2. Material and Methods

### 2.1 Computer Hardware and Software

Lenovo Thinkpad system, with processor properties of 11^th^ generation Intel core i5 @ 2.40 GHz, 8 GB RAM, and 64-bit operating system was used for the computational studies in this study. The software packages used include Marvin sketch for structure drawing and optimization, Autodock tools1.5.7, and Autodock vina developed by The Scripps Research Institute for molecular docking studies, PADEL-descriptor software for molecular descriptor calculations and DTC lab software for 2D-QSAR model development and validation and Microsoft excel 2007 edition.

### 2.2 QSAR analysis

#### 2.2.1 Collection of Dataset and Optimization

A total of 32 synthesized tacrine-cinnamic acid hybrid compounds that were already reported in the literature for their potential to inhibit AChE activity were selected and employed for the development of the 2D-QSAR model (Supplementary Table 1) (Chen et. al., 2017). The inhibitory activity of these compounds was reported to be in the nano molar range and demonstrate the potential of molecules to inhibit 50% AChE enzymatic activity. The IC_50_ values of these 32 molecules were converted to the negative logarithm (PIC_50_) to reduce the skewness in the biological activities. The chemical structures of tacrine-cinnamic acid hybrid compounds and their IC_50_ values are depicted in Table 1. The molecular structures of 32 selected molecules were prepared using Marvin sketch software of Chemaxon and were saved in “mol” format. The optimization of these structures was also performed using the same software using Merck Molecular Force Field 94 (MMFF94).

#### 2.2.2 Molecular descriptor calculations and data normalization

Molecular descriptor calculations and data optimization was performed using PaDEL descriptor software. Briefly, all 32 optimized molecular structures were exported in PaDEL descriptor software for the calculations of molecular descriptors. The PaDEL descriptor software is a product of the Pharmaceutical Development Exploration Laboratory developed by Chun Wei Yap. In the present study, the current version of the software computed a total of 1875 descriptors (1444 1D, 2D descriptors, and 431 3D descriptors), which included topological, electrostatic, autocorrelation, spatial, constitutional, geometrical, and thermodynamic descriptors (Yap, 2011; Banjare et. al., 2021; Chakravarti and Alla, 2019).

#### 2.2.3 Data pretreatment, dataset division, and QSAR model generation

In the present study, we generated a Multiple Linear Regression QSAR model by using a genetic algorithm approach for variable selection. This is a search heuristic method that is used to mimic the process of natural selection. Data pretreatment was used to remove inter-correlated and constant descriptors. The variance cut-off value for constant descriptors and inter-correlation variance was set to 0.01 and 0.85 respectively. Kennard-Stone’s algorithm division technique is a rational approach for dataset division by employing the most diverse molecules in the training set. We performed the division of the test and training sets through this algorithm division technique in the present study and 30% of the selected molecules from the dataset were employed for the test dataset and the remaining 70% of the selected molecules from the dataset were used as the training dataset. We used rational dataset division rather than a random approach to avoid any loss of information due to splitting. All these steps were performed using software DTC-QSAR version 1.0.6 of the DTC lab which is a complete QSAR modeling solution (http://teqip.jdvu.ac.in/QSAR_Tools/) (Yap, 2011; Banjare et. al., 2021; Chakravarti and Alla, 2019).

#### 2.2.4 Internal Validation

We performed the internal validation of the generated QSAR model by employing a crossvalidation technique, which provides sufficient information regarding the reliability of the prediction of the generated model (Yap, 2011; Banjare et. al., 2021; Chakravarti and Alla, 2019). Further, we employed a leave-one-out cross-validation technique for the internal validation of the current model. Cross-validated Q^2^_cv_ was calculated by using the following equation:

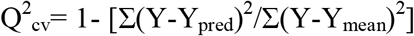

In this equation, Y represents the observed value (PIC_50_), Y_pred_ represents the value predicted by the model, and Y_me_an represents the mean of the observed values of the training set. The squared correlation coefficient (R^2^)value of the training set is a measure of the quality of the developed model, however, it is not much reliable as its value correlates to the number of used descriptors. To overcome this, we calculated the new parameter R^2^_adj_ by using the following equation:

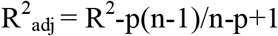

In this equation, p represents the number of descriptors used, and n represents the number of molecules used in the training set for model development. The difference between R^2^ and R^2^_adj_ was less than 0.3, suggesting that the number of descriptors selected is acceptable. We computed all these variables to assess the quality of the developed model internally.

#### 2.2.5 External Validation

To validate the developed model externally and internally, we subjected the dataset divided through Kennard-Stone’s algorithm, along with the selected variables (using a genetic algorithm) and their biological activity values to MLR plus Validation 1.3 tool of DTC lab. The statistical features proposed by Golbraikh and Tropsha (2002) for a robust QSAR model with good predictive potential are depicted in Table 2.

**Table 2:** The statistical features proposed by Golbraikh and Tropsha for a robust QSAR model with good predictive potential.

| S.No. | Parameter | Threshold value |
| --- | --- | --- |
| 1 | $Q^2$ | Threshold value $Q^2 > 0.5$ |
| 2 | $R^2_{\text{train}}$ | Threshold value $R^2_{\text{train}} > 0.6$ |
| 3 | $\vert R^2_0 - R'^2_0\vert $ | Threshold value $\vert r_0^2 - r'^2_0\vert < 0.3$ |
| 4 | k or k' | $0.85 < k < 1.15$ & $0.85 < k' < 1.15$ |
| 5 | $R^2_{\text{test}} - R^2_0 / R^2_{\text{test}}$ | Threshold value $R^2 - R^2_0 / R^2 < 0.1$ |

Where R^2^_0_ depicts the coefficient of squared correlation between the observed and predicted values and R^’2^_0_ depicts the between predicted and observed of the test set.

Also, another error-based judgment of the prediction quality of the test dataset was proposed by Roy et. al. (2016) based on the calculation of Mean Absolute Error (MAE) which was calculated by using the following equation:

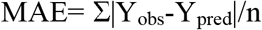

Where Y_obs_ represents the observed value and Y_pred_ represents the predicted value of the test set and n represents the number of compounds in the test set. For a model with good prediction potential, the value of MAE 95% should not be more than 0.15, which is the value of MAE calculated after 5% of high residuals data are omitted from the test data set.

#### 2.2.6 Y randomization test

In this test, we generated the MLR models by keeping the values of descriptors constant and shuffling the value of Y variables. This shuffling could be done n number of times to generate n number of models and the value of R^2^ and Q^2^ was compared for each model with the originally generated models. Ideally, to be a robust QSAR model, the values of R^2^ and Q^2^ of the random models should be as low as possible (Rücker et. al., 2007). We further calculated another parameter in Y randomization,^c^R^2^_P_, by using the following equation:

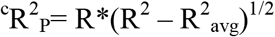

In this equation, R^2^avgdepictsthe average squared correlation coefficient of the random models. For a robust model, the value of ^c^R^2^Pshould ideally is greater than 0.5 (http://teqip.jdvu.ac.in/QSAR_Tools/).

#### 2.2.7 Applicability Domain

Principle 3 suggested by the Organization for Economic Co-operation and Development (OECD), is recommended to define the applicability domain of the developed QSAR model for the prediction of new data points. The applicability domain of a given QSAR model is the chemical structure space in which the model makes predictions with maximum reliability. To obtain the structural and response outliers, the applicability domain approach was used in QSAR model generation (Roy et. al., 2015; Gramatica, 2007). In the current study, two approaches were employed for defining the applicability domain of the generated QSAR model. The standardization approach developed by Roy et. al. (2015) was employed for defining the applicability domain of the generated model. Briefly, 99.7% of the populations of test and training sets lie within the range mean ±3 standard deviation (SD). Therefore, mean ± 3 SDrepresents the zone where most of the training set compounds belong. Any compound that lies outside this zone is considered dissimilar and is considered the outlier in the generated QSAR model.

The second approach that we followed for defining the applicability domain was by plotting the Williams plot from the generated QSAR model. This approach was proposed by Gramatica (2007) based on the leverage value. This approach allows us to predict the position of a new molecule in the acceptable domain of the defined model or outside it. To develop the William plot, we calculated the leverage *h_i_*, of each data point of the test and training dataset according to the following equation:

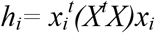

In this equation, *x_i_*, depicts the descriptor vector of the considered data point, X depicts the descriptor matrix, and *X^t^*depicts the transpose of the descriptor matrix. The threshold leverage *h*was* calculated by using the following equation:

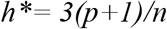

Where p depicts the number of variables and n depicts the number of compounds in the training set. In the present study, a molecule with *h_i_* > *h*\* was considered to be outside the applicability domain, however, it may not be the outlier considering the small standardized residual. These compounds were considered influencers in the QSAR model. A value of ± 3 for standardized residuals was employed as a cut-off value for accepting predictions inside the applicability domain. The leverage and the standardized residual were combined for the characterization of the applicability domain in the present study.

#### 2.2.8 Screening of the herbal molecules

A total of 85 herbal molecules were selected from the literature and were screened for their potential to inhibit AChE activity by using the screening module of the DTC-QSAR v1.0.6 software of the DTC lab. Before the screening, descriptors were calculated for all 85 herbal molecules by using PaDEL-descriptor software and their PIC_50_ values were predicted by using the equation developed from the generated QSAR model.

#### 2.2.9 *In-Silico* Docking study analysis

The *in-silico* docking study was performed using AutoDock Vina (Trott and Olson, 2010). Briefly, optimized structures of the 09 most active predicted herbal compounds were drawn using MarvinSketch software of Chemaxon with the Crystal Structure of Recombinant Human Acetylcholinesterase (PDB ID: 4EY7). The docking simulations were done using AutoDock Vina and visualization of the results was performed using the AutoDock Tools package of The Scripps Research Institute.

#### 2.2.10 Central Nervous System Multiparameter Optimization (CNS MPO) analysis

It is a well-established fact that for any compound to be active in the disorders of the brain, it should cross BBB so that it can impart its beneficial effects. To attain this desirability, Wager et. al. proposed a CNS MPO score for defining the ability of a compound to act centrally. The value of CNS MPO score ranges from 0 to 6 for a molecule and the predicted score should be ≥ 04 to be accepted as a possible CNS candidate (Wager et. al., 2016). CNS MPO score was calculated by employing six ADME properties, which include lipophilicity, calculated partition coefficient (ClogP), calculated distribution coefficient(pH 7.4) (ClogD), molecular weight (MW), topological polar surface area (TPSA), number of hydrogen-bond donors (HBDs), and most basic center (pKa). In our study, the CNS MPO score of all the 9 predicted active compounds was calculated using the MarvinSketch software of ChemAxon.

## 3. Results

### 3.1 In Silico QSAR analysis

In the present study, we employed the genetic algorithm of the descriptor to develop a multiple linear regression (MLR) QSAR model by using DTC-QSAR v1.0.6 of DTC lab by employing a total of 04 descriptors. The MLR equation generated for the model is given below:

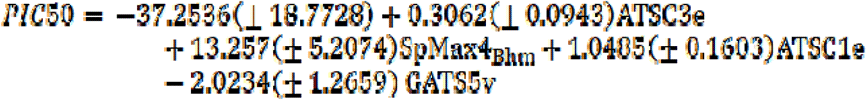

Where, N_train_ = 23, R^2^_train_ = 0.7976, R^2^_adj_ = 0.7526, Standard Error of Estimation (SEE) = 0.1928, Q^2^(LOO) = 0.7061, average r^2^_m_(LOO) = 0.606, delta r^2^_m_ (LOO) = 0.1379, ntest= 9, R^2^test = 0.83852 and MAE (95% data) = 0.13567.

From the above model, the most significant descriptors having positive contributions include ATSC3e, SpMax4_Bhm, and ATSC1e, along with GATS5v that was having negative contribution in the developed model. The details of the 04 descriptors used to generate the above QSAR model are given in Table 3 below.

**Table 3:** Definition of the descriptors used to generate the above QSAR model.

| S. No. | Descriptor | Description | Class | Type | Contribution |
| --- | --- | --- | --- | --- | --- |
| 1 | ATSC3e | Centered Broto-Moreau autocorrelation - lag 3 / weighted by Sanderson electronegativities | Autocorrelation | 2D | Positive |
| 2 | SpMax4_Bhm | The largest absolute eigenvalue of Burden modified matrix - n 4 / weighted by relative mass | Burden Modified Eigen values | 2D | Positive |
| 3 | ATSC1e | Centered Broto-Moreau autocorrelation - lag 1 / weighted by Sanderson electronegativities | Autocorrelation | 2D | Positive |
| 4 | GATS5v | Geary autocorrelation - lag 5 / weighted by van der Waals volumes | Autocorrelation | 2D | Negative |

In the developed model, we observed the correlation coefficient value of 0.79762 and 0.83852 for R^2^train and R^2^test, respectively, suggesting a good extrapolation between the training and test dataset. Further, the robustness of any QSAR model is determined by the difference between the R^2^tram and Q^2^ (< 0.5%). In the QSAR model developed in the present study, we observed a small difference between R^2^_train_ and Q^2^ (< 0.5%), which further validates the robustness of our generated QSAR model. The plot between predicted PIC_50_ and observed PIC_50_ is depicted in Figure 1. It can be observed in the figure that the values of the test set are in close agreement with the training set, suggesting the development of a robust QSAR model.

**Figure 1:**
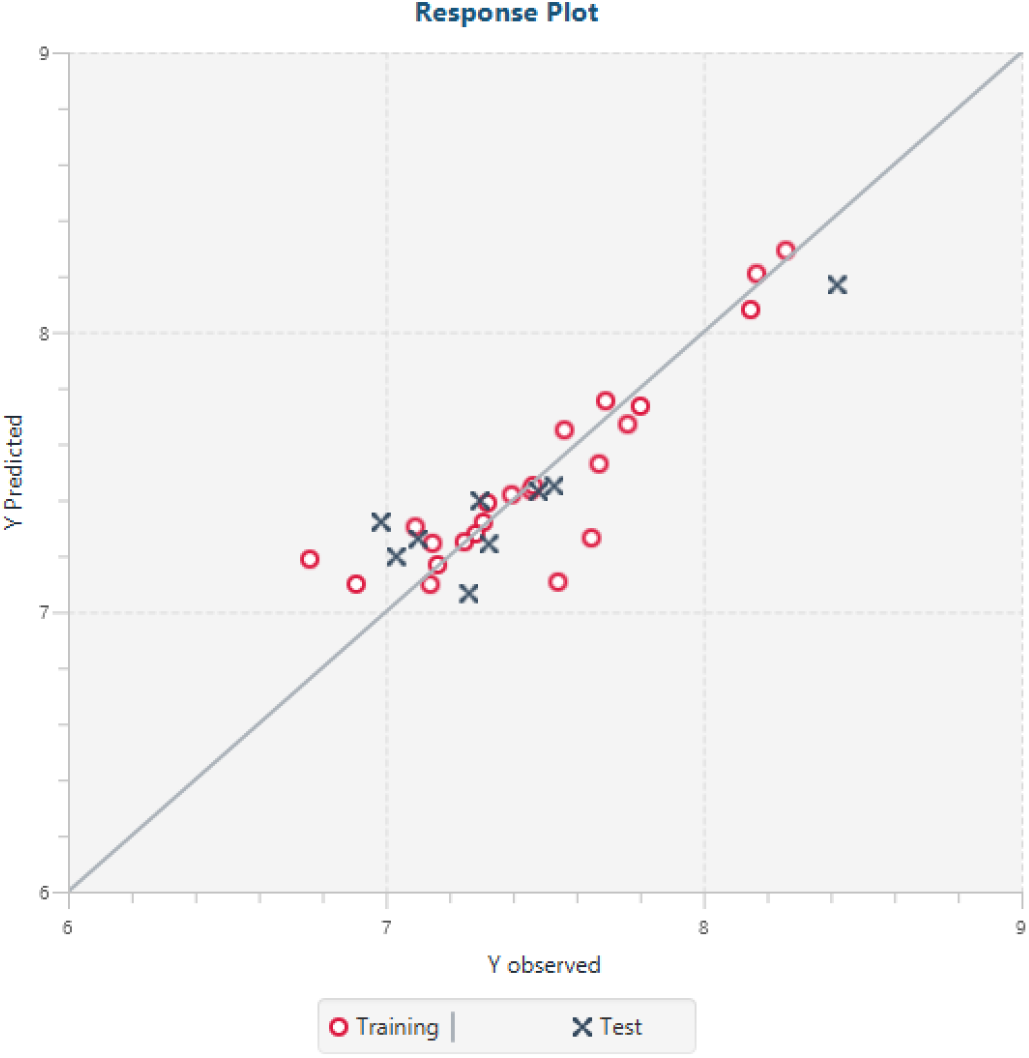
The scatter plot between predicted PIC_50_ & observed PIC_50_ of data points. We performed the Y randomization test on the training set to further establish the robustness of the developed QSAR model. In the present study, we performed 50 Y-random tests and our results suggest that for all the models except one, the values of r^2^ and q^2^ were less than 0.5 (Figure 2). The^c^R^2^_P_value was calculated to be 0.71, which is greater than 0.5, suggesting that the developed QSAR model is robust, rather than derived merely due to chance.

Moreover, in the standardized approach of the Applicability Domain, none of the compounds from the training and test dataset was found to be outside the Applicability Domain, which confirms the good predictability of the developed QSAR model. Further, the scatter plot of standardized residual and leverage, Williams plot, suggests that there is only one compound of the test set having *h_i_* value greater than *h\** (0.682), suggesting it is an influencer and not the outlier as the standardized residual was observed within the limit (Figure 3).

**Figure 2.**
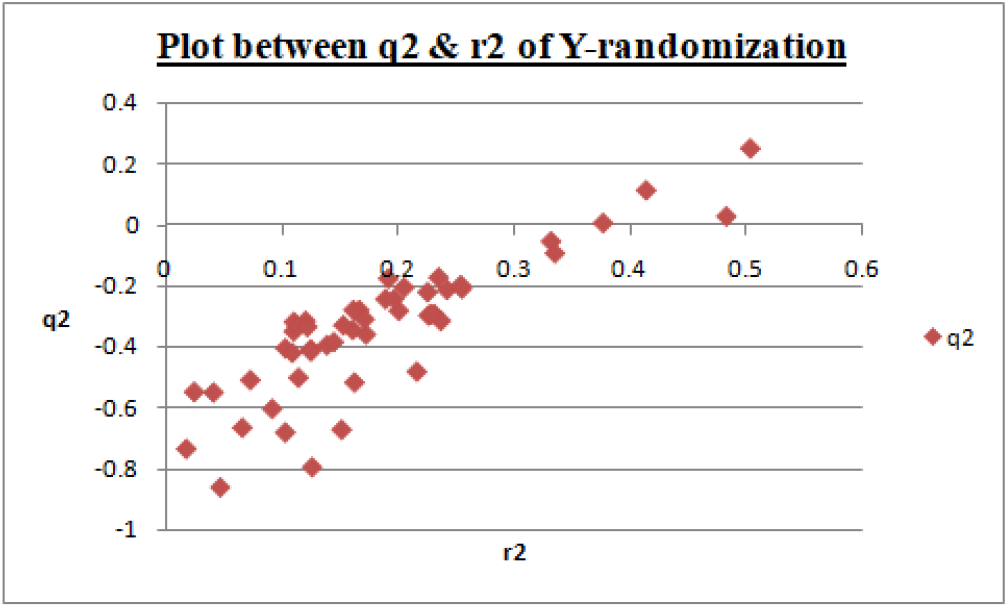
Plot between q^2^ & r^2^ of 50 Y scrambled models.

**Figure 3:**
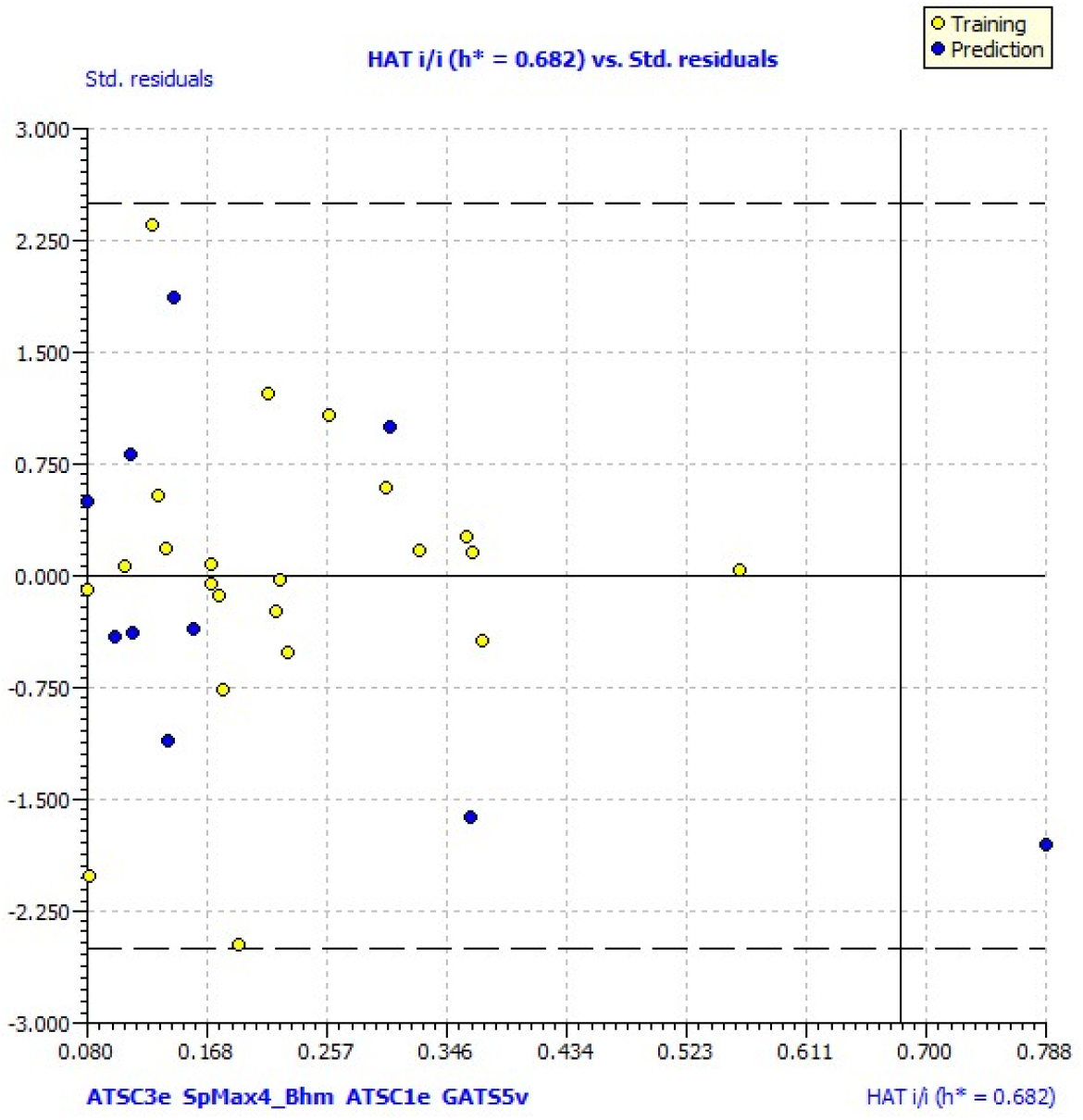
The scatter plot of standardized residuals and leverages (Williams plot)

**Figure 3:**
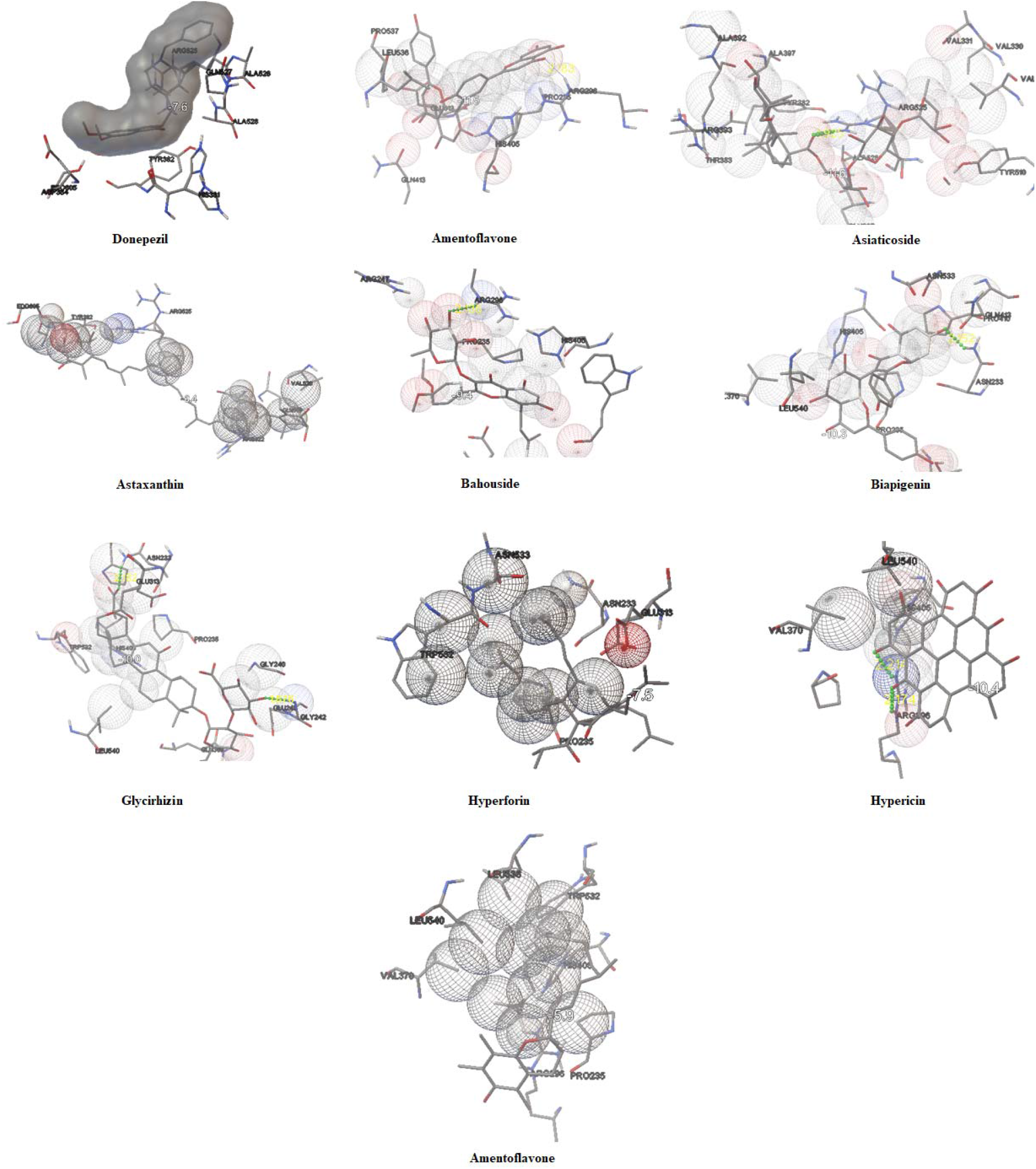
Interaction of donepezil, amentoflavone, asiaticoside, astaxanthin, bahouside, biapigenin, glycyrrhizin, hyperforin, hypericin, and tocopherol with the human AChE.

### 3.2 Virtual Screening, CNS MPO score, and in silico docking analysis

Table 4 depicts the predicted PIC50 of the top 09 herbal compounds with their calculated CNS MPO score. Subjecting herbal molecules to the developed QSAR model suggests amentoflavone, asiaticoside, astaxanthin, bahouside, biapigenin, glycyrrhizin, hyperforin, hypericin, and tocopherol are the most promising herbal molecules for inhibiting AChE activity when compared to donepezil. The PIC_50_ values of these compounds were observed in the range of 6.50 nM to 10.499 nM when compared to donepezil (PIC_50_ = 6.56 nM). Our results suggest that asiaticoside (PIC_50_ = 10.499 nM), glycyrrhizin (PIC_50_ = 10.4 nM), astaxanthin (PIC_50_ = 8.97 nM), biapigenin (PIC_50_ = 7.999 nM), hyperforin (PIC_50_ = 7.61 nM), and amentoflavone (PIC_50_ = 7.377 nM) as the most promising molecules as their PIC50 values were predicted to be considerably lower. Moreover, the CNS MPO score of the herbal molecules was observed in the range of 1 to 3.76, where amentoflavone displayed the best CNS MPO score of 3.76, followed by asiaticoside, biapigenin, and bahouside with the CNS MPO score of 3 each. CNS MPO score is critical for predicting the BBB penetration and CNS activity of the drugs. Considering PIC50 values and CNS MPO score of amentoflavone, our results suggest that amentoflavone could cross the BBB and therefore could be able to inhibit AChE inside CNS.

**Table 4:** Predicted PIC_50_ values and CNS MPO score of the herbal molecules.

| S. No. | Name of the compound | Predicted $PIC_{50}$ values (nM) | CNS MPO score |
| --- | --- | --- | --- |
| 1 | Donepezil | 6.56 | 4.39 |
| 2 | Amentoflavone | 7.377 | 3.76 |
| 3 | Asiaticoside | 10.499 | 3 |
| 4 | Astaxanthin | 8.97 | 2.5 |
| 5 | Bahouside | 7.295 | 3 |
| 6 | Biapigenin | 7.999 | 3 |
| 7 | Glycyrrhizin | 10.4 | 2.93 |
| 8 | Hyperforin | 7.61 | 2.75 |
| 9 | Hypericin | 6.61 | 1 |
| 10 | Tocopherol | 6.50 | 2.73 |

These molecules were then subjected to an *in-silico* docking study by using AutoDock Vina against human AChE (PDB ID: 4EY7) to further consolidate the results and provide proper validation of the predictions made by the developed QSAR model. The results of the molecular docking study are depicted in Table 5 and Figure 4. Our results suggest that amentoflavone, asiaticoside, astaxanthin, bahouside, biapigenin, glycyrrhizin, hyperforin, hypericin, and tocopherol are having good potential to interact with the human AChE. The docking score for these molecules was observed in the range of −5.9 to −11.5 kcal/mol when compared to donepezil (−7.6 kcal/ mol). Our results suggest that ARG 296, ARG 525, and ASN 233 are the major amino acid involved in the ligand-protein interaction. The results of the docking study predicted hypericin and glycyrrhizin as the most promising inhibitor of AChE as indicated by the docking score of −10.4 and −10 kcal/ml along with a total of 02 hydrogen bonds formed with the protein. Likewise, asiaticoside and amentoflavone formed only 01 hydrogen bond with the protein and were having a docking score of −11.6 and −11.5 kcal/mol, respectively, suggesting them to be a promising inhibitors of human AChE.

**Table 5:** Screening of herbal molecules through *in-silico* docking study using human AChE (PDB ID 4EY7)

| S. No. | Name of the compound | Dock Score (kcal/mol) | No. of H-Bond | Amino acid involved | Atoms of the ligand | Bond length (Å<sup>0</sup>) |
| --- | --- | --- | --- | --- | --- | --- |
| 1 | Donepezil | -7.6 | 0 | - | - | - |
| 2 | Amentoflavone | -11.5 | 01 | ARG 296 | O of OH group | 2.183 |
| 3 | Asiaticoside | -11.6 | 01 | ARG 525 | O of OH group | 1.925 |
| 4 | Astaxanthin | -8.4 | Nil | - | - | - |
| 5 | Bahouside | -9.4 | 01 | ARG 296 | O of OH group | 2.135 |
| 6 | Biapigenin | -10.3 | 01 | ASN233 | O of OH group | 2.152 |
| 7 | Glycyrrhizin | -10 | 02 | ASN233<br>GLY242 | O of OH group<br>O of OH group | 2.182<br>1.876 |
| 8 | Hyperforin | -7.5 | 0 | - | - | - |
| 9 | Hypericin | -10.4 | 02 | ARG296<br>ARG296 | O of OH group<br>O of OH group | 2.176<br>2.214 |
| 10 | Tocopherol | -5.9 | 0 | - | - | - |

## 4. Discussion

The development of a statistically significant QSAR model with good predicting ability depends on obtaining descriptors that are the representation of the variations in the structural aspects of the compounds against a specific target (Tropsha, 2007). In the present study, we obtained a total of 1875 descriptors from PaDEL software in Microsoft excel format. The results were further exported to DTC-QSAR v 1.0.6 software for data pre-treatment and normalization. The constant and interrelated descriptors that were having a square correlation coefficient of more than .85 were removed in the data pre-treatment process. These pre-treated data were divided into training and test sets using the Kennard-Stone algorithm, where 70% of the compounds (23 datasets) were placed in the training dataset, and the remaining 30% (9 datasets) were placed under the test dataset. Table 6 provides the details of the acceptability criteria proposed by Golbraikh and Tropsha (2002) and the results suggest that our generated QSAR model passes the Golbraikh and Tropsha criteria of QSAR model robustness. Our results are in line with the previous literature reports where QSAR models have been validated robustness using the criteria suggested by Golbraikh and Tropsha (2002).

**Table 6:** Golbraikh and Tropsha acceptable model criteria.

| S.No. | Parameter | Threshold value | Model Score |
| --- | --- | --- | --- |
| 1 | $Q^2$ | Threshold value $Q^2 > 0.5$ | 0.70613 |
| 2 | $R^2_{\text{test}}$ | Threshold value $R^2_{\text{test}} > 0.6$ | 0.83852 |
| 3 | $\vert R^2_0 - R'^2_0\vert $ | Threshold value $\vert r_0^2 - r'^2_0\vert < 0.3$ | 0.17215 |
| 4 | k or k' | $0.85 < k < 1.15$ & $0.85 < k' < 1.15$ | 0.99892 or 1.00049 |
| 5 | $R^2_{\text{test}} - R^2_0 / R^2_{\text{test}}$ | Threshold value $R^2 - R^2_0 / R^2 < 0.1$ | 0.04138 |

The regression statistics in terms of p-value and t-values are depicted in Table 7. In line with these reports, our results suggest that the coefficients of the descriptors are statistically significant at a 95% confidence interval. Furthermore, previous findings suggest that there should be no correlation between the selected descriptors for an efficient QSAR model against a specific target (Verma et. al., 2017). Table 8 depicts the correlation matrix of all the descriptors used for the development of the QSAR model and it clearly indicates that there is no correlation between the 04 descriptors selected for the model generation.

**Table 7:**
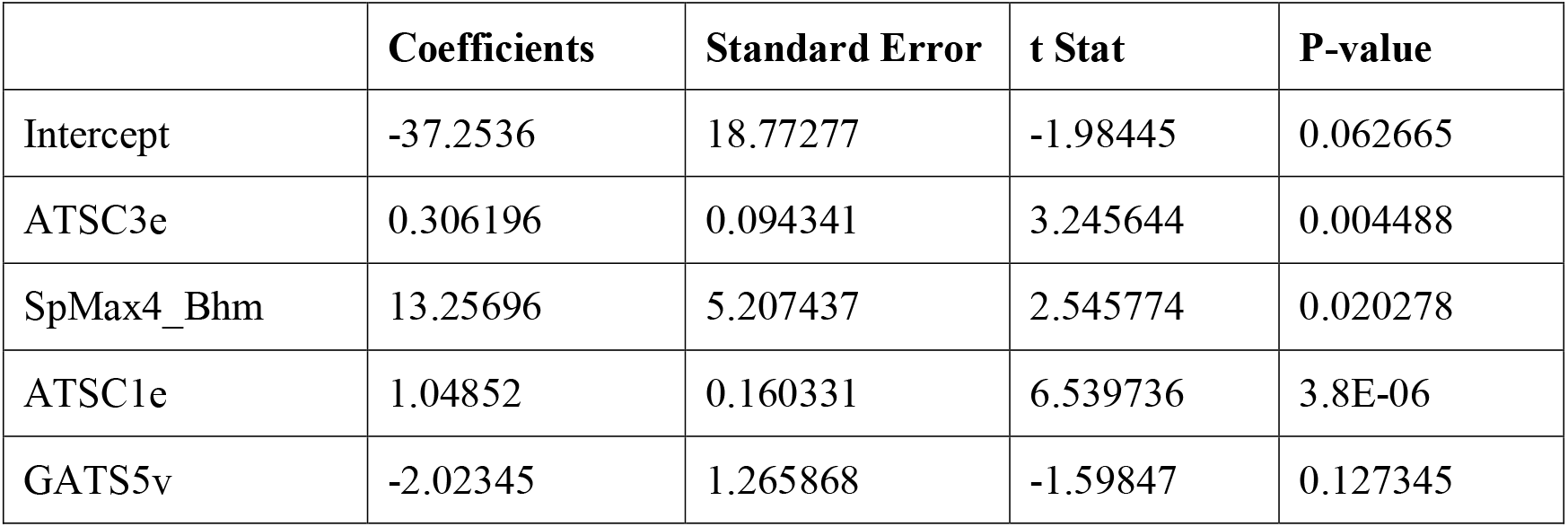
Regression statistics of the descriptors.

**Table 8:** Correlation matrix of the descriptors.

|  | ATSC3e | SpMax4_Bhm | ATSC1e | GATS5v |
| --- | --- | --- | --- | --- |
| ATSC3e | 1 |  |  |  |
| SpMax4_Bhm | 0.124742 | 1 |  |  |
| ATSC1e | -0.06827 | 0.20614 | 1 |  |
| GATS5v | 0.446121 | -0.09939 | 0.010124 | 1 |

The scatter plot between standardized residual versus experimental PIC50 shows an unsystematic scattering of data points above and below the baseline of zero of standard residual, which is a clear indication that there is no systematic error in the developed QSAR model (Figure 5).

**Figure 5:**
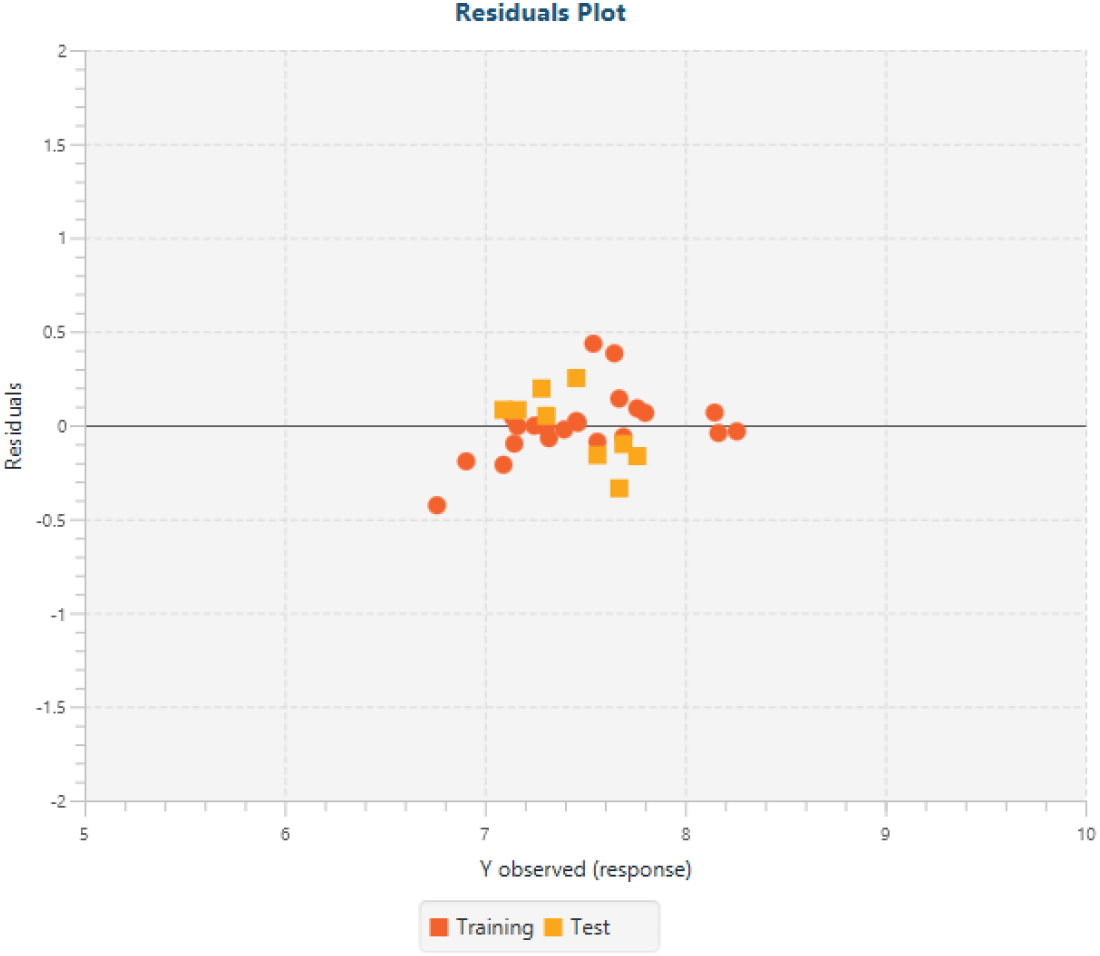
The scatter plot between residuals and experimental PIC_50_ of data points.

The scatter plot of standardized residual and leverage, which is known as the Williams plot, suggest that there is only one compound of the test set having *hi* value greater than *h\** (0.682), suggesting it is an influencer and not the outlier as the standardized residual was observed within the limit (Figure 3). From the literature, an inference can be made that the compounds having a *hi*-value less than *h\** & values of standard residuals between ± 3 are said to be within the applicability domain (Fan et. al., 2018). Williams plot of our developed QSAR model hence clearly indicates its robustness and also that AChE inhibitory activity of the 32 data points is well predicted and is reliable.

A total of 85 herbal compounds selected from the literature were subjected to the developed QSAR model to predict the best potential molecules against AChE (Chen et. al., 2017). Our results predicted tocopherol is the most promising herbal molecule for inhibiting AChE activity when compared to donepezil. These molecules were screened through an *in-silico* docking study to validate our QSAR model to suggest the most promising molecule against human AChE. Molecular docking studies have been used previously to predict the activity of herbal and synthetic molecules against various drug targets, including AChE (Kumar et. al., 2019). Docking study provides us with reliable data regarding the most promising molecules that can have beneficial effects against disorders like AD (Kumar et. al., 2019), diabetes (Zargari et. al., 2018), depression (Dong et. al., 2021), Parkinsonism (Mirza et. al., 2015), etc. the docking results predicted hypericin (forming two hydrogen bonds with the protein), glycyrrhizin (forming two hydrogen bonds with the protein), asiaticoside (forming one hydrogen bonds with the protein) and amentoflavone (forming one hydrogen bonds with the protein) as the most promising inhibitor of AChE as indicated by the docking score of −10.4, −10, −11.6 and −11.5 kcal/mol, respectively. The docking score and receptor interaction of these molecules was better than that observed for donepezil, suggesting that these molecules could be a promising AChE inhibitor. Accumulated evidence in the recent past also suggests that hypericin, glycyrrhizin, asiaticoside, and amentoflavone are having the potential to impart beneficial effects during neurodegenerative conditions (Béjaoui et. al., 2017; Song et. al., 2013; Hossain et. al., 2015; Rong et. al., 2019). Moreover, these molecules have been reported to have good antioxidant and anti-inflammatory potential (Rizk et. al., 2021; Oh et. al., 2013; Dellafiora et. al., 2018; Pradeep et. al. 2019; Frattaruolo et. al., 2019; Nurlaily et. al., 2012). Considering the fact that oxidative stress and inflammatory stress play a major role in the pathogenesis of AD and other neurodegenerative disorders, these molecules could attenuate neurodegeneration inflicted during oxidative and inflammatory stress. Furthermore, there have been reports where these molecules have been reported to have inhibitory effects against AChE. Therefore, the results of the docking study not only predicted the most promising molecules against human AChE, but also validated the predictability, robustness, accuracy, and reliability of the developed QSAR model.

It is evident from the literature that to date hundreds of compounds have been predicted through docking studies to have AChE inhibitory activity, however, very few of them reproduced the results in the pre-clinical studies. This is probably due to the fact that these molecules either are having poor bioavailability or fail to cross the BBB, which is a major limitation in achieving the beneficial effects against CNS disorders like AD, especially with herbal molecules (Pardridge, 2012). Therefore, to address this we calculated the CNS MPO score to predict the potential of herbal molecules to cross the BBB. Interestingly, the QSAR and molecular docking study predicted hypericin, glycyrrhizin, asiaticoside, and amentoflavone as the most promising molecules against AChE, but only amentoflavone possessed the desired CNS MPO (3.76) score which was close to 04 suggesting this molecule can efficiently cross the BBB. All other remaining 08 herbal molecules possessed CNS MPO scores less than or equal to 03 indicating their limitation to cross the BBB which is a prerequisite for any drug to act centrally. The results of the QSAR model, docking study, and CNS MPO suggest that amentoflavone is the most promising molecule that could have a beneficial effect during AD by targeting the AChE enzyme within the CNS.

## 5. Conclusion

In the present study, we attempted to develop a robust and reliable QSAR model to predict the AChE inhibitory activity of herbal compounds and the results were validated through docking studies. The QSAR model developed was found robust and statistically accepted with good predictability based on the statistics metrics, and therefore can be used efficiently to screen synthetic and herbal molecules for their potential against human AChE enzyme. The molecular docking studies further validated our prediction based on the QSAR model with most of the herbal compounds possessing better docking scores than the standard. We concluded hypericin, glycyrrhizin, asiaticoside, and amentoflavone as the most promising molecules against human AChE. However, results of the CNS MPO score suggest that amentoflavone may have good bioavailability and could efficiently cross BBB to impart its beneficial effects within the brain. In overall conclusion, we predict and suggest amentoflavone to be the most promising molecule against AD with the proposed mechanism of AChE inhibition within the CNS.

## 7. Acknowledgement

The authors acknowledge Govt. College of Pharmacy, Rohru for providing the facilities to conduct this study.

## 8. Author’s Contribution

AT contributed to designing the study, performed QSAR and docking studies, and prepared the first draft of the manuscript. AP, VS, and AK contributed to the study desigining, technical inputs, and manuscript editing. VM contributed to designing the study, molecular docking, manuscript preparation, and editing analysis of the results.

