## Supplemental Table 1 for "2D-QSAR and Molecular Docking based virtual screening of the herbal molecules against Alzheimer’s Disorder: An approach to predict CNS activity"

**Supplementary Table 1:** A total of 32 synthesized tacrine-cinnamic acid hybrid compounds selected and employed for the development of the 2D-QSAR model (Chen et. al., 2017)

| 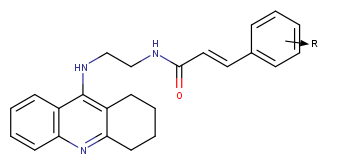 | | | |
| --- | --- | --- | --- |
| Name Of Compound | R | IC_50_ value (nM) | PIC_50_ |
| 1 | H | 92.5 | 7.033 |
| 2 | 2-CH_3_ | 80.6 | 7.093 |
| 3 | 3-CH_3_ | 78.9 | 7.102 |
| 4 | 4-CH_3_ | 34.3 | 7.464 |
| 5 | 2-OCH_3_ | 123.8 | 6.907 |
| 6 | 3-OCH_3_ | 47.4 | 7.324 |
| 7 | 4-OCH_3_ | 22.5 | 7.647 |
| 8 | 2,3-diOCH_3_ | 47.8 | 7.32 |
| 9 | 2,5-diOCH_3_ | 72.3 | 7.14 |
| 10 | 2,3,4-triOCH_3_ | 17.3 | 7.761 |
| 11 | 3,4,5-triOCH_3_ | 20.3 | 7.692 |
| 12 | 3,4-OCH_2_O- | 49.3 | 7.307 |
| 13 | 2-Cl | 50.4 | 7.297 |
| 14 | 3-Cl | 34.9 | 7.457 |
| 15 | 4-Cl | 21.3 | 7.671 |
| 16 | 4-F | 68.6 | 7.163 |
| 17 | 4-Br | 27.4 | 7.562 |
| 18 | 2-CF_3_ | 52.1 | 7.283 |
| 19 | 4-CF_3_ | 33 | 7.481 |
| 20 | 2-NO_2_ | 7.1 | 8.148 |
| 21 | 3-NO_2_ | 3.8 | 8.42 |
| 22 | 4-NO_2_ | 6.8 | 8.167 |
| 23 | 4-Cl, 3-NO_2_ | 5.5 | 8.259 |
| 24 | 3-OCF_3_ | 56.5 | 7.247 |
| 25 | 4-Methyl Carbonate | 71.2 | 7.147 |
| 26 | 2-OBn | 103.2 | 6.985 |
| 27 | 3-OBn | 40.1 | 7.396 |
| 28 | 4-OBn | 29.5 | 7.53 |
| 29 | 3-OMe, 4-OBn | 15.8 | 7.801 |
| 30 | 2-NH_2_ | 28.7 | 7.542 |
| 31 | 3-NH_2_ | 54.7 | 7.262 |
| 32 | 4-NH_2_ | 173.3 | 6.761 |
